# Striatal dopamine supports reward reactivity and learning: A simultaneous PET/fMRI study

**DOI:** 10.1101/2020.06.24.169722

**Authors:** Finnegan J Calabro, David Montez, Bart Larsen, Charles Laymon, William Foran, Michael Hallquist, Julie Price, Beatriz Luna

## Abstract

Converging evidence from both human neuroimaging and animal studies has supported a model of mesolimbic processing in computing prediction errors, which form the basis of reward learning behaviors. However, direct evidence demonstrating how human dopamine signaling in the basal ganglia contributes to learning has been hampered by limitations of individual imaging modalities. Here, we present data from a large (N=81, 18-30 year olds), multi-modal neuroimaging study using simultaneously acquired task fMRI, affording temporal resolution of reward system function, and PET imaging with [^11^C]Raclopride (RAC) assessing striatal D2/3 receptor binding. Results indicated that task-related dopamine release in the ventral striatum, measured as a change in RAC binding, was greater among those who demonstrated successful reward learning on a probabilistic map learning task. This learning response was specific to the ventral striatum and was not present in fMRI BOLD reward response activation. This provides support for considering task-related DA release in ventral striatum as a key signal for translating reward outcomes into a learning signal, rather the representing the reward outcome in isolation. These data provide novel, human *in vivo* evidence that dopaminergic function may support reward reactivity as well as reward learning as distinct processes.

## 1. Introduction

The functional roles of striatal dopamine (DA) signaling have been increasingly studied, due to their wide-ranging contributions to learning, motivational, and motor processes. Converging evidence from human neuroimaging studies, including Positron Emission Tomography (PET) and functional magnetic resonance imaging (fMRI), as well as electrophysiology, voltammetry, and optogenetic studies, provide compelling evidence that ventral striatal dopamine signals the value of reward outcomes relative to their expectation (1). Critically, this signaling has been found to support updating (i.e., learning) in addition to reward reactivity (2) but their relative contribution is not well understood.

Work using animal models has characterized important contributions of DAergic processes in supporting unique aspects of reward processing and learning. Phasic activity of DA neurons in the ventral tegmental area (VTA) signals the difference between reward receipt and the expected value of that reward (2,3), termed reward prediction error (RPE) (4). More generally, such responses are generated upon the updating of expectation of total future rewards as part of the temporal difference (TD) model (5,6). By this model, any information that causes a revision of future expected rewards generates an error signal, providing the basis for reward learning behaviors. Recent work has begun to dissociate the contributions of VTA and nucleus accumbens (NAcc) to RPE, with VTA identifying the presence of a reward, and NAcc DA release reflecting reward expectation (7).

Critically, encoding RPEs in this manner enables DAergic processes to play a key role in learning reward contingencies. Striatal DA neurons modulate long term potentiation (LTP) and depression (LTD) of synaptic strength (8,9) (for review see (10)). Dopaminergic activation has further been shown to regulate dendritic spine growth (11), providing mechanisms by which RPEs can affect synaptic plasticity, potentially enabling the updating of future reward expectations.

Neuroimaging studies in humans have shown a similar role for ventral striatal DA mechanisms in reward processing. Prediction error responses have been demonstrated in NAcc responses in fMRI studies (12–14), and a growing body of PET studies have supported the view that these can be linked directly to DAergic processes (15,16). Interestingly, reward related ventral striatal (VS) BOLD activity has been found to extinguish when subjects view rewards passively (17) and related PET-derived dopamine responses are small when received rewards are not necessary for optimizing future performance (e.g., learning) (18). In contrast, VS engagement has been reported when rewards are successfully used to learn reward contingencies, that is, during successful reward learning (19). Recent work has suggested that VS RPE responses may directly support learning by adaptively coding the PE as scaled by the variance of the distribution (20). These studies point to a role for striatal DA in which responses scale with prediction error, but depend critically on the use of these signals in active learning processes.

However, the relationships between DAergic signaling and neuronal activation, and the relative contribution of these processes to reward reactivity and reward learning, is still not fully understood in humans. While PET can provide a direct measure of different DA processes, measurements occur over extended periods of time (e.g., at the whole session level) limiting our ability to assess what aspects of learning are associated with DA. fMRI can be used to assess trial, epoch, and condition specific responses to rewards and learning but it does not provide a direct measure of DA. Here we used multimodal, molecular MRI (mMRI) approach to simultaneously obtain raclopride-based PET measures of D2/D3 receptor function and fMRI trial level BOLD response during a reward learning task that assesses both reward receipt and learning processes. By simultaneously acquiring these measures as subjects perform a reward learning task, we are able to characterize how these distinct measures of underlying neural activity contribute to computational reward learning mechanisms, both in terms of trial-to-trial activation responses, and in the contribution of DA-specific signaling mechanisms. Our results support a model of DA signaling in which reward responses are encoded in ventral striatal regions in relationship to their contribution to ongoing reward learning processes. Further, we demonstrate that epoch and condition specific neuronal activation responses do not predict successful learning, nor do they fully account for the learning-specific DA signaling. Our results provide direct evidence in behaving human subjects that distinct aspects of DA physiology and function support active learning processes in addition to reward reactivity that can also inform impaired function such as in psychopathology (addiction, psychosis, mood disorders).

## 2. Results

### 2.1. Behavioral Performance

Performance on the task was assessed both within-task, based on the proportion of moves made to the higher probability map location and by RL model fit, as well as post-task, based on the map learning assessment. RL model fits exhibited a bimodal distribution (see Supplemental Figure S2), supporting a distinction between “learners” and “non-learners”, which we define as those who were (ΔAIC>10 compared to a “guessing” model) and were not well fit (ΔAIC<10) by the RL model. Overall, this produced two similarly sized groups, with 41 participants were classified as learners and 39 as non-learners (see Supplemental Table S1 for subject characteristics). These groups did not differ on age (learners 23.6+/-3.2, non-learners 22.9+/-4.1, p=0.37), gender (learners 49% female, non-learners 54% female, p=0.66), or IQ (learners 108.5+/-9.4, non-learners 108.8+/-10.6, p>0.9). As expected, learners showed a greater proportion of optimal moves (defined as choices to the higher true probability location) than nonlearners overall (T=3.3, χ^2^=11.1, p=0.0008, see Figure 1A). Not surprisingly, this difference was not present in the first block (T=0.57, p=0.57), but emerged beginning with the second block (T=2.36, p=0.02) as subjects had time to explore the map. Since making optimal responses is highly associated with the ability of the RL model to fit the data, this dissociation is somewhat ensured by the group definition. However, there was also a significant difference in post-task map learning performance, with learners able to identify 77.2% of map pairs correctly, while nonlearners identified 62.3% of pairs correctly (F=50.5, p<10^-9^, see Figure 1B). Since failing to be well fit by an RL model does not guarantee that participants did not learn, this provides an independently characterized validation of the description of the groups as those who did and did not learn the reward contingencies. Finally, both groups showed increasingly faster responses with later blocks (T=-6.64, χ^2^=99.2, p<0.0001), but there was no significant difference between groups (T=0.095, χ^2^=0.019, p=0.88), nor was there a significant group by block interaction (T= −0.79, χ^2^=0.62, p=0.42, see Figure 1C).

**Figure 1.**
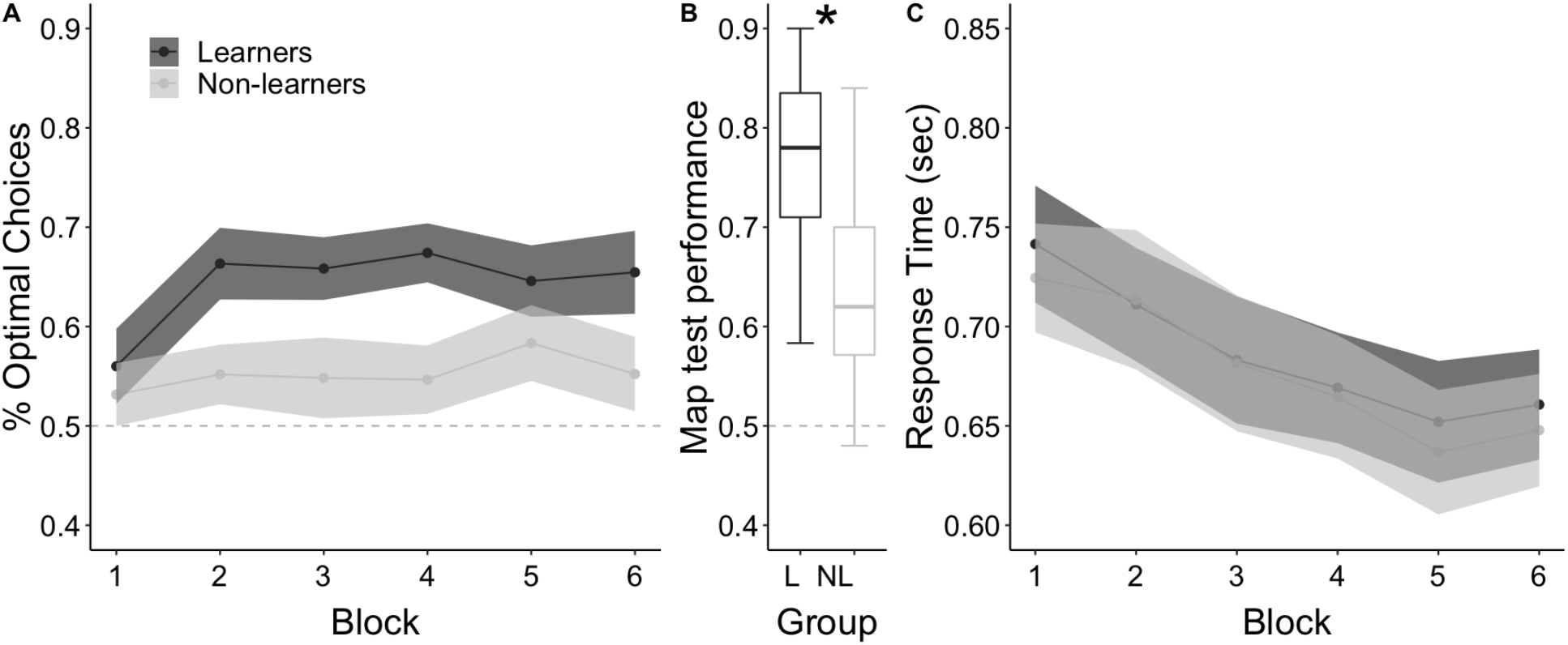
Task performance by learner group. Performance as quantified by (A) proportion of optimal trials by blocks, (B) distribution of post-task assessments for learners (L) and non-learners (NL), and (C) response time by block, split by learner category. Dashed horizontal line indicates chance performance.

### 2.2. BOLD reward response

fMRI data was analyzed by computing the contrast between rewarded and non-rewarded trials. We found significant activation across the striatum, as well as an extensive cortical network. A contrast between rewarded and non-rewarded (“null”) trials revealed a large cluster with distinct peaks centered on the bilateral ventral striatum (see Figure 2), extending upward along the dorsal thalamus, and downward into the parahippocampal gyrus. Additionally, there was a large medial cluster along the posterior and mid cingulate cortex, a cluster in ventromedial PFC (vmPFC) extending into the rostral cingulate, two sets of bilateral inferior frontal gyrus clusters (BAs 44 & 46), bilateral posterior-to-middle insula clusters, and several occipital and visual association clusters (See Supplemental Figure S3 and Supplemental Table S2 for cortical regions).

**Figure 2.**
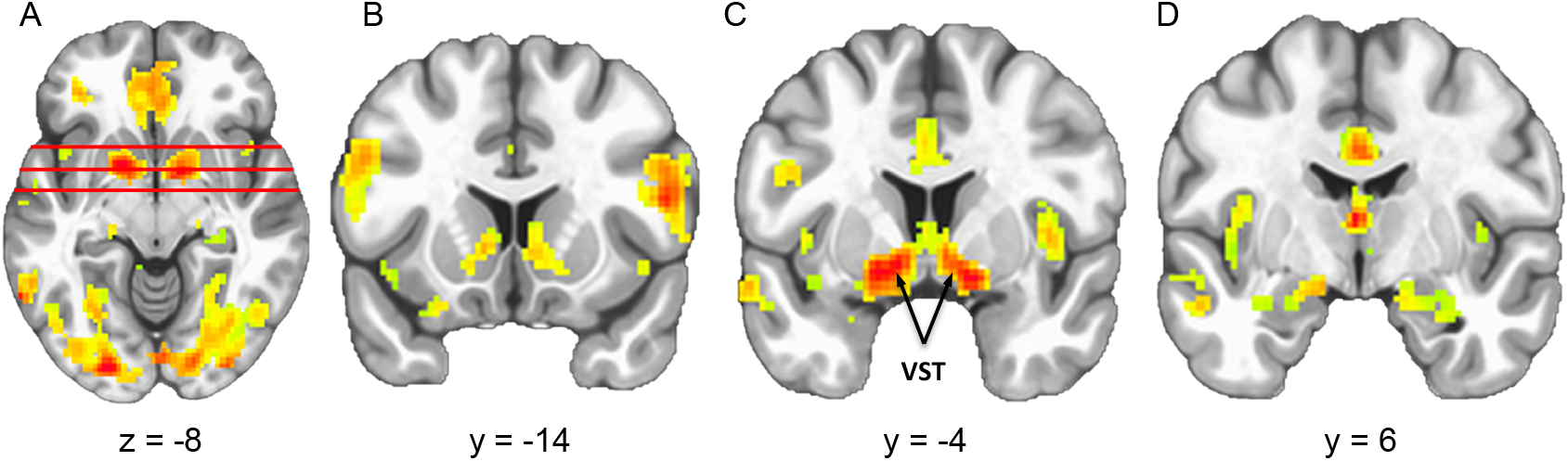
Cluster corrected task fMRI activation for the contrast of rewarded vs. non-rewarded trials, analyzed for the reward receipt epoch. Three coronal slices (depicted on the axial slice, left) are shown to illustrate extent of striatal activation, which was centered on the ventral striatum (VS, panel C). See Supplemental Table S2 for clusters of activation, and Supplemental Figure S3 for whole brain activation maps.

### 2.3. Striatal DA release

PET data was analyzed to assess dopamine (DA) release over the entire task period (approximately 30min). A decreased binding potential compared to baseline indicates fewer available receptors, suggesting the release and binding of DA which displaces the tracer. We assessed DA release voxelwise across the entire striatum to identify anatomical regions which had significant changes in their [^11^C]RAC BPnd. We found four significant clusters which survived cluster correction (family wise p<0.01) of decreased BPnd: bilateral clusters centered on the ventral striatum (VST) regions extending slightly into the precommisural caudate nucleus (CN), and bilateral clusters contained within the precommisural dorsal putamen (PUT) (see Figure 3).

**Figure 3.**
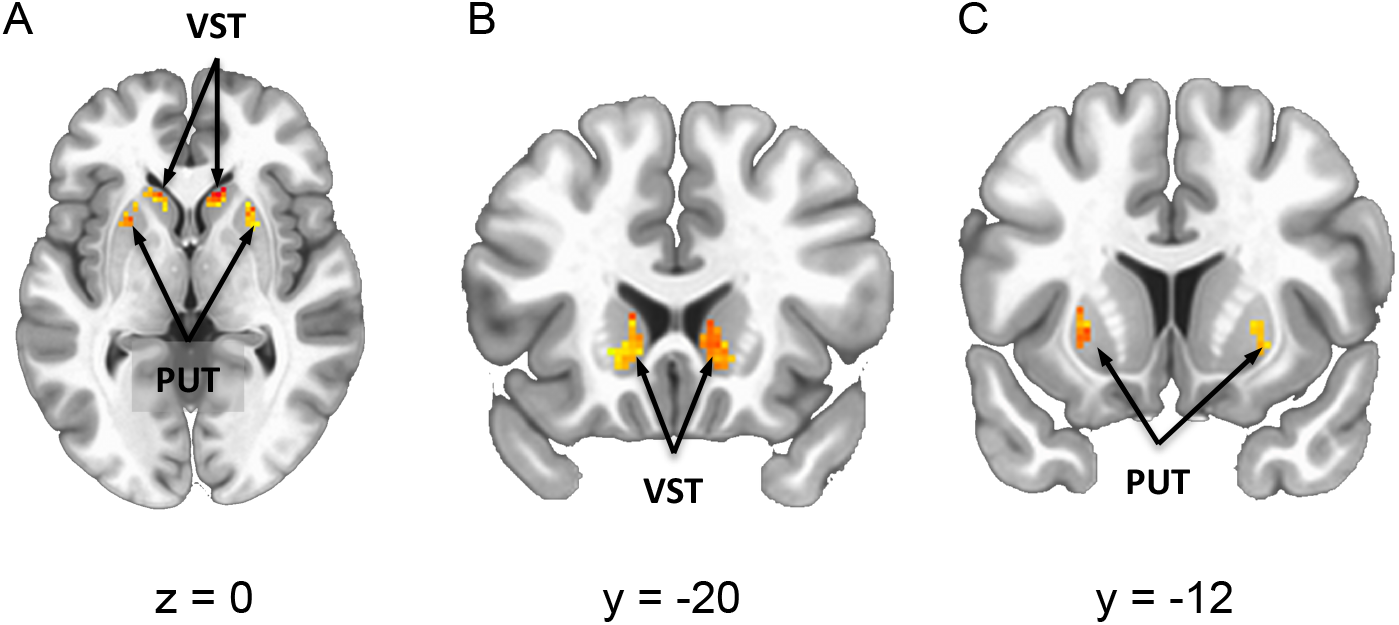
Cluster corrected maps of significant task-related change in RAC BP. Slices are shown at (A) z=0 (axial) (B) y = −20 (coronal) (C) y = −12 (coronal). Four clusters of activation were observed (left slice), including a bilateral set focused primarily on the ventral striatum (VS, panel B), and a bilateral set focused on the precommisural dorsal putamen (PUT, panel C).

### 2.4. DA release and BOLD activation by learners and non-learners

To assess the relationship between DA and successful reward learning on our task, we compared RAC task effects between learners and non-learners. We focused on anatomically-defined regions that contained significant overall task effects in the RAC data, namely, the VST, CN and PUT, as defined above. This indicated that task-related changes in RAC BP in the VST (F=5.3, p=0.025), but not CN (F=0.14, p=0.71) or PUT (F=0.27, p=0.61) were significantly greater among learners compared to non-learners (see Figure 4B), while no difference was observed in baseline binding potential between groups (VST: F=1.58, p=0.21; CN F=1.53, p=0.22; PUT F=0.14, p=0.71, see Figure 4A). In contrast, aggregate BOLD activation (reward vs neutral trials, combined across all reward probability levels) did not differ between groups for any of the three regions (VST: F=0..29, p=0.59; CN F=0.07, p=0.80; PUT F=0.11, p=0.31). A similar pattern of results was observed when separating BOLD responses by reward level (low/medium/high, see Supplemental Material).

**Figure 4.**
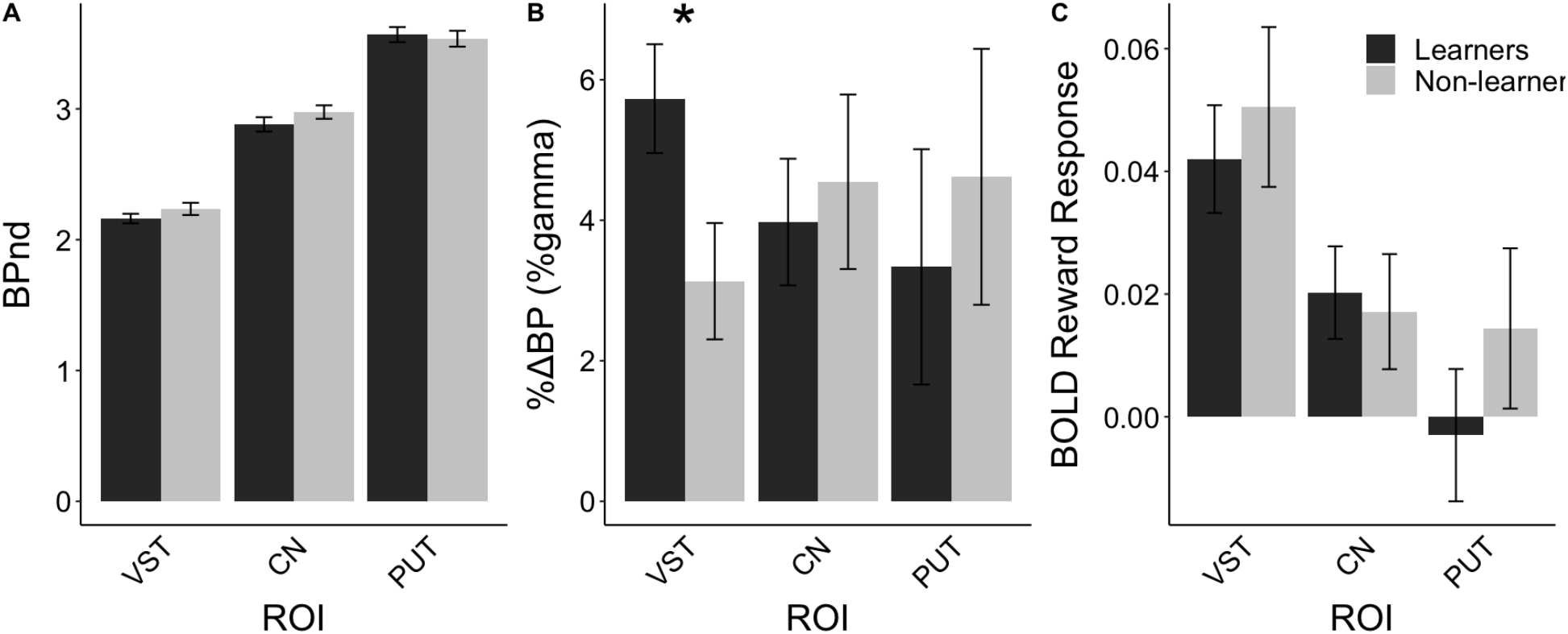
Comparison of (A) baseline RAC BPND, (B) task-related DA release, as measured by a change in RAC BP, and (C) BOLD reward response comparing all rewarded to all non-rewarded trials. Responses are averaged across voxels within each of three striatal regions (VST: ventral striatum, CN: precommisural caudate, PUT: precommisural dorsal putamen) for those who did (“learners”) and did not (“non-learners”) demonstrate reward learning behavior as assessed by the reinforcement learning model.

### 2.5. Association of DA and with reward learning behaviors

To further characterize the association of dopamine release with reward learning performance, we compared the relationship between task-related change in RAC BP with individual model parameters based on the consensus RL model. We found a significant association between overall model evidence (indicating support for performance consistent with the RL model, relative to a guessing model) and DA release in the VST as indexed by %ΔBP (T=3.4, p=0.0012, see Figure 5A). This supports the association (indicated above) between learners and non-learners, and the existence of a continuous relationship between these variables suggests that it is unlikely to be due to the choice of classification threshold. Furthermore, we found a significant quadratic relationship between learning rate for positive outcomes and %ΔBP (linear term, T=2.72, p=0.0085; mean-centered quadratic term, T=-2.38, p=0.021, see Figure 5C), such that high DA release was observed for those with learning rates near 0.5; learning rates that were either very low or very high were associated with diminished DA release. We did not find any significant relationship between DA release and either the softmax temperature parameter (beta, T=1.08, p=0.28) or the negative outcome learning rate (T=0.04, p=0.96). No such associations were found between RL parameters and aggregate BOLD responses (see Supplemental Figure S6).

**Figure 5.**
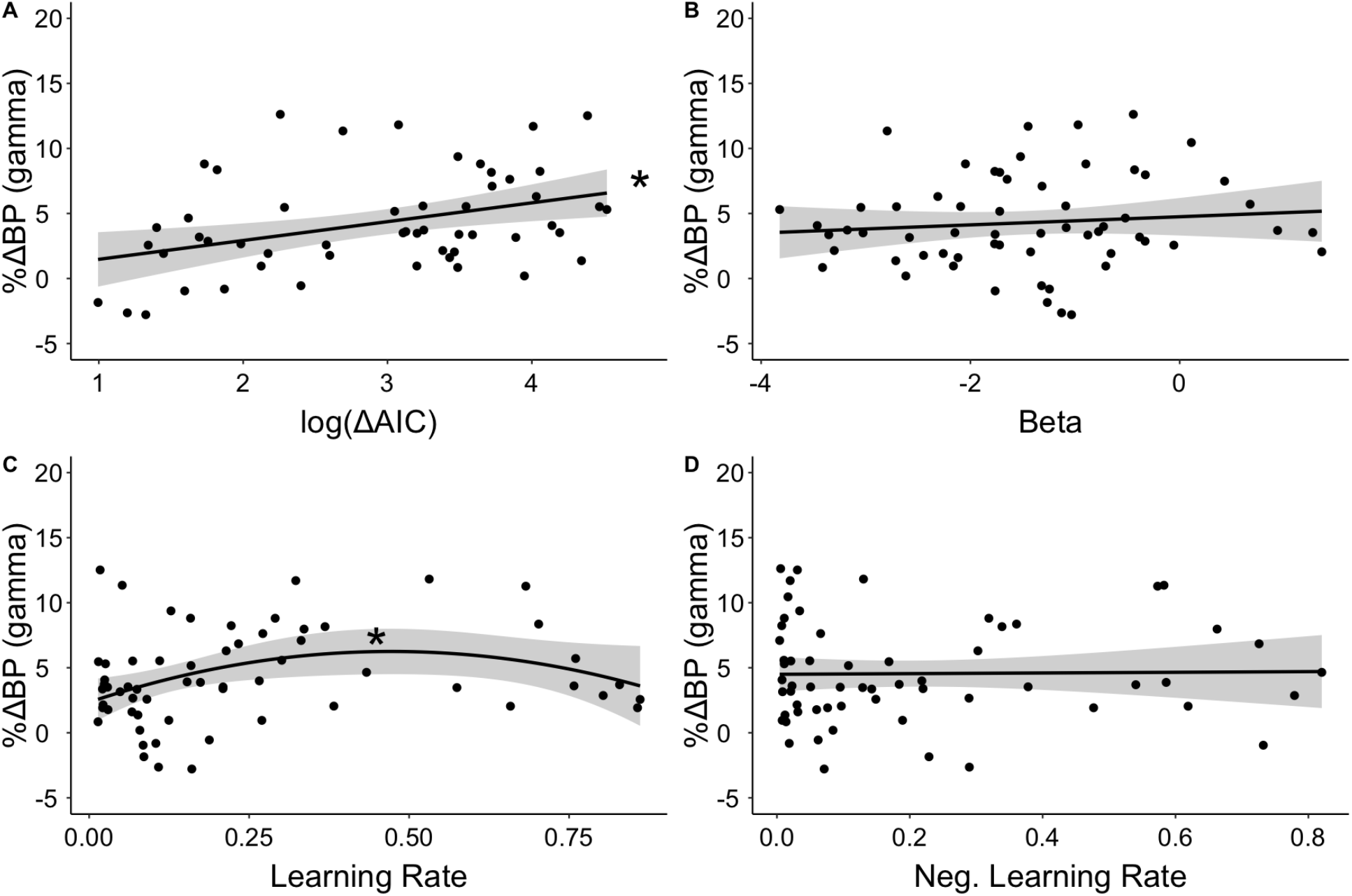
Association of task-related DA release in the VST and RL model parameters, including (A) overall model evidence relative to a guessing model, (B) softmax temperature parameter, (C) learning rate for positive PE trials, and (D) learning rate for negative PE trials.

## 3. Discussion

To our knowledge, this study provides the first direct evidence from simultaneous PET-fMRI imaging for dopaminergic contributions to reward learning processes. We leveraged simultaneous PET-fMRI acquisition during a probabilistic reward learning task to characterize the role of DA. Methodologically, this approach is valuable for linking temporally sensitive fMRI measures to transmitter-specific PET responses. Since learning is a dynamic process and may vary significantly upon repeated attempts, obtaining this information within a single scan session provides an unprecedented view of the functional neurophysiological processes underlying reward learning. This unique dataset revealed that performing a reward learning task elicited both a BOLD activation as well as PET DA response in the ventral striatum. We leveraged a comparison between those with successful (“learners”) and unsuccessful (“non-learners”) performance on our probabilistic reward learning task to identify neurophysiological processes associated with reward learning. These results indicated that while both groups showed a similar overall BOLD response to reward outcomes in the VST, they were distinguished by PET-assessed, task-related DA response, which was significantly greater among learners and was correlated RL model fit parameters. This pattern of effects suggests that DA responses during reward learning tasks do not passively reflect reward receipt but are instead also part of an active learning process in which participants translate unexpected reward outcomes into task-specific knowledge.

Striatal DA has been associated with a number of distinct functions, including motor processes, reward response, and learning (21). Dissociating these contributions has been difficult, especially in human studies, in part because these functions often co-occur. The distinction of learner and non-learner sub-groups in our study allows us the opportunity to characterize their relative contribution. First, since the sensorimotor aspects of the task were matched across all participants, it is relatively unlikely that these could directly contribute to the group differences we observed. In addition, in the fMRI BOLD analyses, we contrasted rewarded and non-rewarded trials. There was no difference in the motor aspects of these tasks, since the classification of rewarded or non-rewarded did not occur until the reward is presented, which was after the completion of the motor response. Thus, motor differences are unlikely to contribute to the BOLD responses presented here. Furthermore, we matched the reward schedule across participants, such that all participants received the same pattern of reward feedback and with matched timing, meaning that visuomotor differences are unlikely to contribute to the group DA effects we have reported.

Separating the contributions of reward receipt and reward learning is even more challenging, since these are often inextricably linked. Work from rodent models has long indicated the reward prediction errors drive DAergic activity in the ventral striatum (22,23), paralleling fMRI studies (12–14,20). Prediction errors can be viewed as both a reward response (magnitude of reward relative to expectation), and as a learning signal since they are the basis for updating in reinforcement learning models. It is, in principle, possible that the group differences in DA that we have reported, could arise due either to a difference in prediction error magnitude, or to the role of DA in translating prediction errors into updated expected value signals. Since our non-learner group was defined as those who could not be well fit by a reinforcement learning model, we do not have reliable trial-by-trial markers of their expectation or, therefore, prediction error. However, it is reasonable to assume that prediction errors should be smaller among learners, since the learning they exhibit should drive expectations to more closely match true expected value. If PEs alone were driving DA responses, we would predict a greater DA response among non-learners since they would fail to reduce PEs through learning. However, since learners showed greater DA responses with no difference in BOLD fMRI responses at the moment of reward receipt, this is unlikely to explain the group differences in DA we have observed. Thus, it is less likely that the group differences in DA reflect differences in the overall magnitude of the prediction error responses.

Instead, we believe these results indicate a role for DA signaling related to learning processes beyond encoding PEs alone. By consideration of the fMRI data, it is unlikely that the PE alone drives group differences in DA release, suggesting that the differences may arise during another part of the learning process, such as the updating of expectations based on reward feedback. Since this process is not time-locked to any specific event with our trial, it is difficult to measure with fMRI, whereas the slow, session-long PET measures would incorporate these processes regardless of when they occur. Another possibility is that the PET measures incorporate ongoing, tonic DA activity, which is not well modeled by the fMRI analysis design. Tonic DA has been proposed to represent vigor of task responses (24–27), which could reflect overall greater task engagement and motivation among our learner group compared to nonlearners. This explanation seems somewhat less likely given that non-learners responded with similar response times as learners, though we did not design the task to test for this directly.

In sum, we suggest that these data support a DAergic contribution to learning, that is, through learned certainty of reward contingencies or through the updating of expected value and reduction in its associated uncertainty based on reward outcomes. This role is supported by associations between DA release and learning rate similar to prior “inverted-U” models (28,29), which posit a sweet-spot wherein either too much or too little DA is related to reduced cognitive performance. In our task, both very low learning rates (which suggest very little information is extracted and/or retained from the reward outcome), and very high learning rates (which implies large updates, such that only the most recent trials contribute to decisions) are detrimental to maximizing rewards. Importantly, both such outcomes were associated with diminished ventral striatal DA responses, whereas peak DA responses were found in individuals with more optimal learning rates. Further work is still needed to tease apart the specific aspects of learning mechanisms.

There are several important methodological considerations when drawing these inferences. First, although a number of PET studies have proposed modeling approaches for single-session task designs, such approaches are still relatively less common. A particular concern is that these models may systematically overestimate pre-task binding potential, or misestimate other parameters (e.g., k2’) of the PET compartment model, creating false positive task effects. We think this is unlikely to account for our results for several reasons. First, we began our task relatively late in the session (35-40min), as indicated by prior simulation studies (30) and relatively conservative compared to other recent approaches (31), so that pre-task BPnd estimates are unlikely to be significantly biased. Second, while any such biases could undermine the main effect of task, they are unlikely to explain the group differences we have seen, since both learners & non-learners began the task at the same time within their scan sessions. Finally, while a mis-estimation of reference tissue model parameters, such as k2’ (the reference tissue rate constant), would affect all voxels, making it less likely that we would see effects constrained to specific clusters within the striatum. Although continued efforts are needed to more fully define optimal modeling approaches, such approaches have the potential to greatly expand our ability to characterize the contribution of DA to behavioral task performance.

These results have important implications to understanding the role of dopamine in reward contexts. Dopamine is still often discussed purely in terms of its role in reward reactivity. However, our data provide *in vivo* simultaneous PET and fMRI evidence supporting a growing body of literature suggesting that reward receipt alone is not sufficient to account for ventral striatal DA responsiveness (e.g., (17)). There are important implications to considering the role of DA in learning beyond reward reactivity. Clinical conditions with DA dysfunction that show abnormal reward processing such as in Parkinson’s Disease (32) may reflect effects on motivated learning in addition or instead of reward reactivity. Adolescent peaks in sensation seeking (33) are frequently deemed to be underlied by elevated reward reactivity (34–36), but these same changes in neurophysiology may impact unique aspects of learning at this time. Our results provide compelling new evidence *in vivo* in humans for multiple roles for dopaminergic function in reward reactivity and learning that can inform comprehensive models of motivation and impact our understanding of lifespan dopaminergic development, and clinical dysfunctions.

## 4. Material and methods

### 4.1. Participants

Eighty-one participants (ages 18-30, mean age 23.3 +/- 3.6, 41 female) completed the full testing protocol, which included a behavioral session and a combined MRI/PET session that was performed on a separate day subsequent to the behavioral session. Participants were recruited from the local population. Exclusions included: major psychiatric illness affecting themselves or a first degree relative; prior neurological illness or injury including loss of consciousness; clinical syndrome levels as assessed by the Adult Self Report (ASR) scale; pregnancy (assessed by urine test), lactation; drug use within the last month; history of alcohol abuse; or contraindications to both MRI (e.g., metal in body) and PET (e.g., prior recent radiation exposure). Participants were consented for both the behavioral and imaging components of the study, and research protocols were approved by the University of Pittsburgh institutional review board, including the radiation safety committee and radioactive drug research committee.

### 4.2. Behavioral Task

Participants performed six 5-minute blocks of a reward learning task while in the scanner. In the task, participants were represented by a frog avatar on a 3×3 map. The map consisted of a grid superimposed on three landmark images (pond, tree, mountain; see Figure 6). Each map location was pre-determined to have a randomized reward probability of 20%, 50%, or 80%. Three grid locations were assigned to each probability level, and the map layout was chosen to avoid common patterns (e.g., three high reward squares in a row). The underlying probabilities were maintained throughout the six task runs. Participants were instructed to explore the ‘map’ to find the cells with most rewards thus encouraging them to learn probability contingencies in order to maximize their earnings (up to $25) over the course of the entire session.

**Figure 6.**
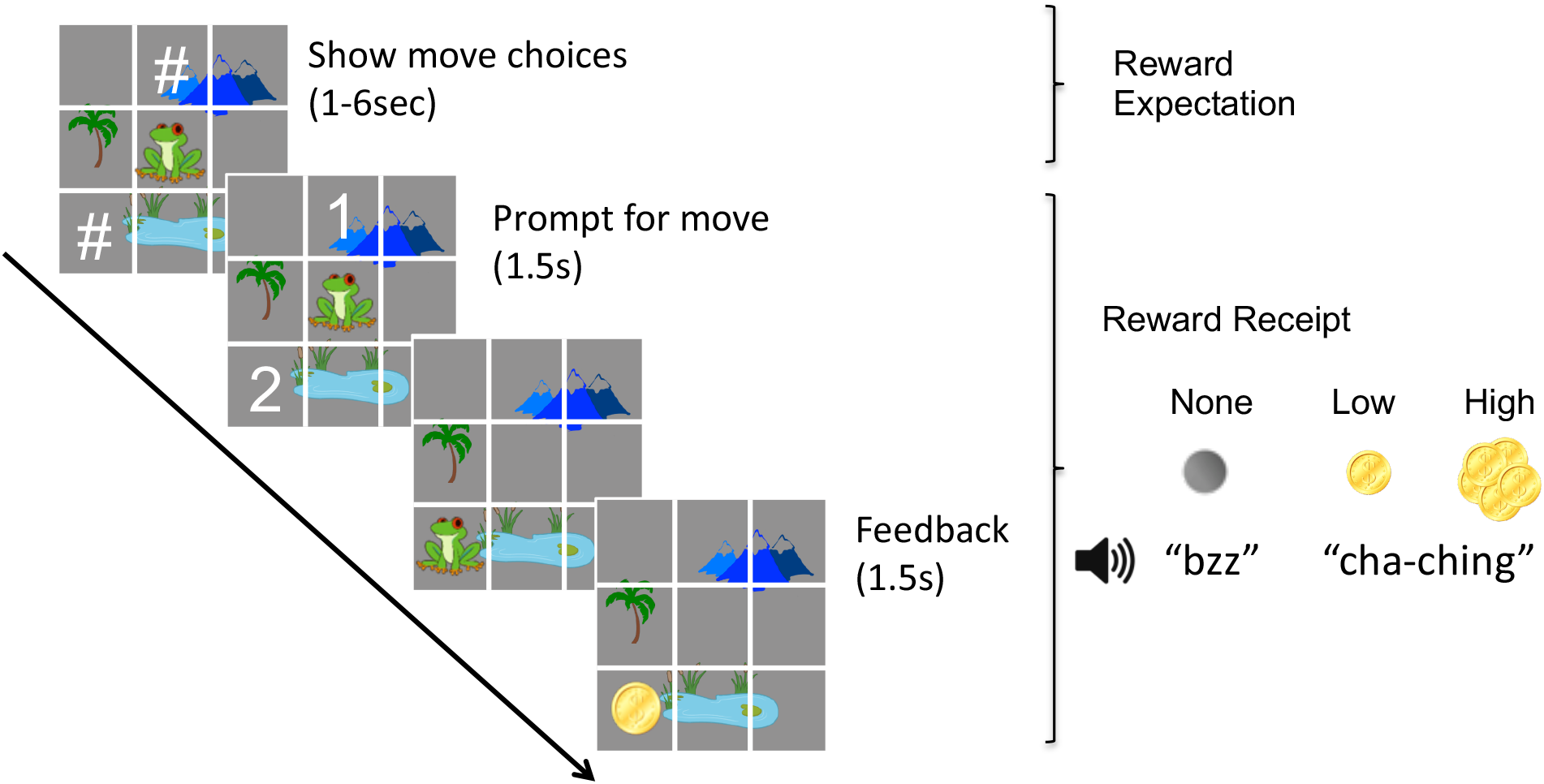
Task schematic. On each trial, subjects were presented two map locations as possible movement choices, indicated by hash marks (“#”). Following a randomized delay, these symbols were replaced by the numbers “1” and “2” such that the subject could make a button press response. Once they responded, the map updated to show their move, and feedback was given as both visual (black circle, single coin, pile of coins) and auditory (flat tone, “cha-ching” sound) feedback to indicate whether a reward was received, and whether it was small (low reward) or large (high reward).

On each trial, participants were presented the option to move to one of two different map locations. Unbeknownst to participants, the two possible locations were pseudo-randomly picked to ensure that the actual reward sequence was matched across participants. This was done so that all participants earned the maximum number of rewards, allowing for equivalent potential for DA involvement, and that all locations were explored. The two locations are initially represented by hash marks (“#”) for a random duration from 1-6 sec, during which subjects were able to consider which square they would move to (thus providing decision making time, and the opportunity for reward expectation), but did not have the cue for response. After this, the symbols changed to display “1” and “2” so that participants could make their response. After selecting a response, the participant’s avatar moved to reflect their new location, and feedback was presented both visually and via auditory cues to indicate whether they received a reward (“cha-ching!”) or not (buzzer), based on the location’s predetermined reward probability, and if so, whether it was a low reward (single coin, 75% of rewarded trials) or high reward (pile of coins, 25% of rewarded trials).

At the end of each block, participants were presented feedback on their total earnings (in arbitrary unit points) thus far. Participants were not given direct feedback about monetary earnings until after the scan was completed. At the completion of the six blocks, participants were tested on their ability to have learned where the most profitable cells were located. In this post scan test, participants were presented 50 pairs of map locations and were instructed to report which of the two squares was better. No feedback was given during this phase, allowing it to be used as a pure measure of the degree to which participants learned the map (that is, there was no exploratory value, as during the main task).

On a portion of trials, after receiving feedback participants performed either a rewarded or unrewarded anti-saccade task. Although this data was modeled in fMRI analyses described below, results are not presented here, as this part was designed to probe cognitive processes for a separate study. In order to learn the mechanics of the task, participants were trained on the task using an abbreviated (single 5-min block) version of the task both during the behavioral session which occurred prior to the scan day, and immediately prior to entering the scanner. Participants were instructed that the maps used during these training sessions would be different than the map used during the scan.

### 4.3. Reinforcement learning model

To assess performance during the task, we fit a reinforcement learning (RL) model to each participant’s trial-by-trial responses. The model tracked the expected value (EV) of each map location following the subject’s moves (*S*), and updated the EV on each trial following feedback via the learning rule

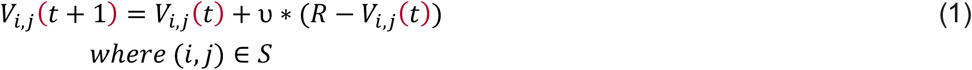

where *V* is the expected value of location (*i,j*), *t* is the trial number, **ν** is the learning rate and *R* is the reward received on the current trial (0 for no reward, 0.5, for a low reward, and 1 for a high reward; low reward values were determined based on the group mean parameter estimate from a preliminary model using a free parameter bounded by 0,1). To assess model likelihood, probability that the subject would choose location “1” was modeled via the softmax function, as

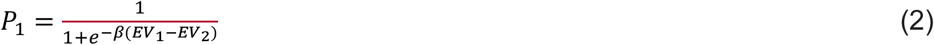

where *β* is the temperature parameter, and EV_1_ and EV_2_ are the expected values of the two map choices for the given trial.

Within this model framework, we considered a number of additional parameters based on a Variational Bayesian Analysis (VBA) using the VBA toolbox in MATLAB (37), as described in the Supplemental Material. The consensus model incorporated separate learning rates for positive and negative prediction error outcomes, similar to previous reports(38). Overall model likelihood was computed as the cumulative probability of the choices made across all trials. Free parameters were determined for each subject by minimizing the overall log likelihood of the model.

### 4.4. MR Data Acquisition

MRI and PET data were collected simultaneously over 90 min on a 3T Siemens Biograph molecular Magnetic Resonance (mMR) PET/MRI scanner. Participants’ heads were immobilized using pillows placed inside the head coils, and participants were fitted with earbuds for auditory feedback and to minimize scanner noise. A 12-channel head coil was used. Structural images were acquired using a T1 weighted magnetization-prepared rapid gradient-echo (MPRAGE) sequence (TR, 2300 ms; echo time [TE] = 2.98 ms; flip angle, 9°; inversion time [TI] = 900 ms, voxel size = 1.0×1.0×1.0 mm). Functional images were acquired using a blood oxygen level dependent (BOLD) signal from an echoplanar sequence (TR, 1500 ms; TE, 30 ms; flip angle, 50°; voxel size, 2.3×2.3 mm in-plane resolution) with contiguous 2.3mm-thick slices aligned to maximally cover the whole brain.

### 4.5. MR Data Analysis

Structural MRI data was preprocessed to extract the brain from the skull, and warped to the MNI standard brain using both linear (FLIRT) and non-linear (FNIRT) transformations. Task fMRI images were processed using a pipeline designed to minimize the effects of head motion (39), including 4D slice-timing and head motion correction, wavelet despiking (40), co-registration to the structural image and non-linear warping to MNI space, local spatial smoothing with a 5mm Gaussian kernel based on the SUSAN algorithm (41), intensity normalization, and high pass filtering *(f* > 0.0125 Hz). Frame-wise motion estimates that were computed, and volumes containing frame-wise displacement (FD) > 0.9mm or DVARS (a measure of total brain signal change) > 21 were excluded from analyses.

First level analysis was performed by modeling all trial events in AFNI’s 3dDeconvolve (42). Events were defined for the hash mark interval (decision making and expectation, modeled as a TENT response persisting up to 24 sec after the event onset), the feedback event (in which participants made their selection and received feedback, modeled with a canonical generalized additive model (GAM) hemodynamic response function), and the preparatory and outcome phases of the anti-saccade task (also modeled with GAM HRFs). Feedback epochs were modeled separately for no reward, low reward, and high reward trials. A 2^nd^ order polynomial regressor was applied separately for each run to account for baseline shifts and drift. In a separate analysis, both expectation and reward receipt phases of the task were modeled separately based on the reward probability level of the selected map location (20/50/80% reward probability).

A contrast was defined between rewarded and non-rewarded trials, weighted by the relative proportion (50% no reward, 37.5% low reward, 12.5% high reward). Group average and group difference activation maps were computed using AFNI’s 3dTtest++. Cluster correction was performed by AFNI’s 3dClustSim program based on permutation testing, using a voxel-wise threshold of *p* < 0.001 and a cluster-wise significance level of **α** < 0.01. For our primary contrast of interest reported below (reward vs. non-rewarded trials), this corresponded to *n* ≥ 96 contiguous voxels.

### 4.6. PET Data Acquisition and modeling

The PET acquisition methods and modeling approach have been previously reported(43). Briefly, we used a 90 minute bolus+infusion administration of [11C]Raclopride (RAC) consisting of a 33-40 mCi dose. Subjects were at rest for the first 35-40 minutes of the scan, at which point they began the reward learning task. We used a modified SRTM model with a cerebellar reference region to quantify both baseline BPnd (binding potential prior to performing the task), as well as the change in BPnd during the task, modeled as a step function. The quantity *γ* by which the BPnd decreased during the task is taken as a measure of DA release, since a reduction in BPnd reflects increase DA occupancy. Details of both the PET acquisition and modeling are reported in detail in the Supplemental Material.

### 4.7. Region of interest selection

To facilitate comparisons between PET and fMRI data, we used regional averages extracted from anatomically defined regions of interest (ROIs). This was done to avoid circularity which would arise from defining regions based on one modality or the other. Anatomical regions were extracted from an atlas defined based on structural T1 and PET ([11C]PHNO) data (44). From this atlas, we considered bilateral ventral striatum, pre/postcommisural caudate, and pre/postcommisural dorsal and ventral putamen.

## 5. Acknowledgements

This work was supported by NIH grant MH080243 to BL. We would like to thank Evan Morris and Shuo Wang for help in designing the PET paradigm and analysis approach. Data collection was expertly performed by Julia Lecht, Matt Misar, Jen Fedor, Jess Graves, and Laurie Thompson. We are grateful to personnel in both the Magnetic Resonance Research Center (MRRC) and the PET center at UPMC Presbyterian, especially James Ruszkiewicz, Tae Kim, Chan Moon, Brian Lopresti, and Hoby Hetherington, for their valuable assistance in designing, implementing, and performing multi-modal imaging acquisitions. We thank the University of Pittsburgh Clinical and Translational Science Institute (CTSI) for their support in recruiting participants. Portions of the PET data presented in this study have been previously reported in Larsen et al, 2020.

