## Supplemental Material for "Striatal dopamine supports reward reactivity and learning: A simultaneous PET/fMRI study"

**Supplemental Methods**

*PET Acquisition*

[11C]Raclopride (RAC) was administered over 90 minutes using a bolus+infusion (B+I) paradigm (1). Participants received a total injected dose of 33-40 mCi with high specific activity. The tracer was prepared in a 60 ml solution with 32.5 ml injected as the initial bolus over a period of two minutes. The remainder was injected at a constant rate of 18.57/h ml over the next 88 minutes (kbol=105 min). PET data were acquired over 90 minutes commencing at the start of tracer injection. Images for attenuation and scatter correction (i.e. images) were developed from the MPRAGE MR images acquired during the PET/MR scan using a combined segmentation- and atlas-based approach (2). RAC data were reconstructed using filtered back projection (30x3-min time frames). The PET images were inspected for interframe motion, and if necessary, corrected using the image fusion tool of PMOD. Data were then aligned to the subject’s MPRAGE, and the same non-linear warp coefficients derived from the MPRAGE were applied to transform data to MNI space.

*PET Modeling*

The 90min RAC acquisition included two parts: [1] an initial, pre-task portion while subjects were at rest, and [2] a later portion beginning 35-40 minutes into the scan during which subjects performed the reward task described above. BP estimates were obtained voxelwise, by fitting the entire time course and including a delta-BP (γ) term to account for changes in BP due to the task, based on previous models to a modified version of the simplified reference tissue model (SRTM) as implemented in MIAKAT (3). Based on previous models (4–7), we used modified version of the SRTM to include a time-varying non-displaceable binding potential (BPnd) term. BPnd was modeled as a step function; that is, BPnd had a constant value pre-task, which decreased by a quantity (*γ*) at the moment the task began and remained at that level for the remainder of the task. The sign of this effect was defined such that a positive γ represented a decrease in BPnd, as we would expect based on an increase in DA release (which decreases the available receptors available for RAC binding and hence BPnd). Specifically, our approach (which has previously been described in (8) stemmed from a re-arrangement of the terms of eq. 7 from (9) given R_1­_=*k*_1_/*k*_2_ and *k*_1_/*k*_2_= *k*_1_’/*k*_2_’, and which produced

$C\left( t \right)=R_{1}C'\left( t \right)+R_{1}\left[ k_{2}'-\frac{R_{1}k_{2}'}{1+BP} \right]C'\left( t \right)\bigotimes e^{\left( \frac{-R_{1}k_{2}'t}{1+BP} \right)}$	(3)

In order to adapt this model to admit a time varying binding potential it may be re-expressed as an equivalent temporal update rule rather than a convolution, which produces

$C\left( t \right)=R_{1}C'\left( t \right)+R_{1}\left[ k_{2}'-\frac{R_{1}k_{2}'}{1+BP} \right]W(t)$	(4)

Where

$W\left( t \right)= e^{\left( -BP \right)} W\left( t-1 \right)+\left[ \frac{1-e^{\left( -BP \right)}}{BP} \right]C'\left( t \right)$	(5)

This temporal update model is equivalent to the current MIAKAT implementation. Our modified implementation replaced$BP$ with $BP\left( t \right)$, modeled as a Heaviside step function in the following way.

$BP\left( t \right)={BP}_{ND}-\gamma h(t)$	(6)

$h\left( t \right)=\left\{ \begin{aligned} 0 (t\leq t_{onset}) \\ 1 (t>t_{onset}) \end{aligned} \right.$	(7)

In this expanded model, C(t) and C’(t) represent the time activity curves of the voxel of interest and reference region respectively, while R_1_ is the ratio of tracer delivery (*k*_1_/*k*_1_’) and *k*_2_’ is the reference region clearance rate constant. Modeling BP(*t*) as a Heaviside step function (eqs. 6,7) allows for time-dependent binding potential having a constant pre-task value BP_ND_, which decreases by a fixed quantity, γ, at the moment the task began (*t_onset_*), and remains at that level for the remainder of the acquisition. Time activity curves (TAC) were extracted for each voxel in the striatum, as well as a regional average TAC derived from the cerebellar gray matter excluding the vermis which was used as the reference region. Least squares fits were performed using Matlab’s lsqnonlin function to estimate four free parameters (R_1_, *k*_2_’, BP_ND_, γ), of which the output variable of interest was the baseline (pre-task) BP_ND_. Observed BP_ND_ values were comparable to prior literature (10–15).

Voxelwise analyses to identify regions with significant BP change were performed using a one sample, 2-sided t-test as implemented by 3dttest++ in AFNI, masked to include the entire striatum. A cluster simulation was performed within this test to identify significant clusters (family-wise p<0.01) given a voxel threshold of p<0.005. Mean head motion (framewise displacement, FD) as computed from the task fMRI data was included as a covariate, though only four voxels had a significant association with motion (at an FDR-corrected p<0.05), and no clusters reached our family-wise error threshold.

*RL Model Specification*

To determine the optimal model specification, we employed a Variational Bayesian Analysis (VBA) using the VBA toolbox in MATLAB (16) comparing a number of competing implementations within the main RL framework. These included three classes of models, in which (1) learning rates were modeled separately for rewarded and non-rewarded outcomes (17), (2) EV decayed towards the initial value on every trial for unvisited map locations (18), and (3) map locations provided additional EV as an “information seeking” term which followed a variable rate decaying exponential based on the number of visits to the given location. We also considered framework parameters, including whether initial EVs were assigned to 0, the mean reward outcome, or were left as free parameters, and whether parameters should vary from block to block. For each subject, we also computed a guessing model to provide a baseline model estimate, in which each trial choice was assigned a 50% probability.

To determine the most parsimonious model across our subject population, we fit every model to each subject’s trial-by-trial choice pattern, then computed group likelihood of each model based on Bayesian Model Comparison (BMC). Within this framework, we first performed a series of family-wise comparisons across models to determine the effects of model parameters and implementations. This provided strong support for modeling parameters across the full session rather than within each block (posterior probability 0.75; model exceedance 1.0), but did not unambiguously differentiate among EV initialization parameterizations (all model exceedances <0.5). Comparing among the model classes provided strong support for the two-learning rate model, in which positive and negative outcomes were associated with separate learning rates (exceedance probability = 0.9944), compared to the base, decay, and information seeking model, as well as models containing combinations of these terms. To confirm these findings and resolve ambiguities in how to model the initial conditions, we next performed the BMC iteratively. On the first iterations, all models (n=41) were compared, and the model with the lowest exceedance probability was dropped from consideration. This procedure was repeated iteratively until a single model remained (see Supplemental Figure S1). This approach confirmed support for the two-learning rate model in which positive and negative outcome trials used separate learning rates. The preferred model had an EV initialization of 0 for each map location, and used static parameters across the task session.

**Supplemental Results**

To further interrogate BOLD VS reward responses, we separated activation into epoch and condition-specific responses. During the expectation phase of the task, overall VS responses were observed as BOLD deactivations. Supplemental Figure S4A summarizes the full timecourses (as shown in Supplemental Figure S5) based on the peak deactivations. A repeated measures ANOVA with main effects of TR, expectation level and learner type, as well as all interactions, was performed on the full timecourses. This produced a significant main effect of TR (**χ**^2^=477.6, p<0.0001), indicating an overall task response, as well as significant main effects of expectation level (**χ**^2^=9.7, p=0.007), seen as a less negative response as reward probability increased, a significant TR*learner type interaction (**χ**^2^=59.9, p<0.001), seen as a shallower, but prolonged, deactivation in non-learners, and an expectation*learner type interaction (**χ**^2^=14.4, p=0.0007), seen as a greater modulation of activation due to expectation level among learners compared to non-learners. During reward receipt, when separated by probability level of the selected map location, a repeated measures ANOVA indicated no overall effect of learner type (F=1.8, p=0.17), nor an interaction between learner type and expectation level (F=1.5, p=0.23, see Supplemental Figure S4B).

**Supplemental Table S1.** Participant summary, split by learner/non-learner classification. Means and standard deviations (in parentheses), as well as Odds Ratios (OR) between groups, and p-values from a two-sample T-test are reported for each measure.

**Supplemental Table S2.** Regions showing greater reward activation compared to no reward trials.

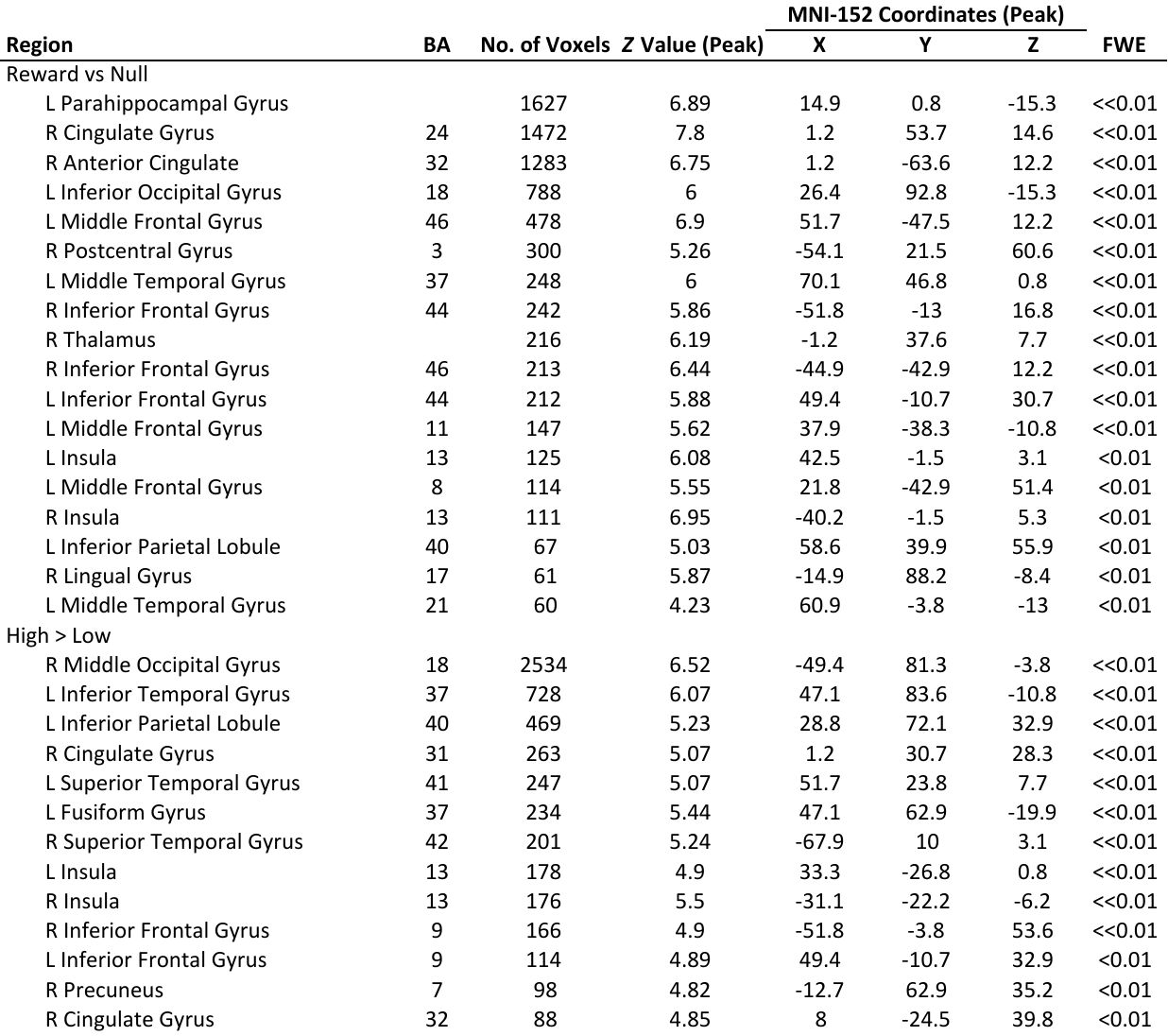

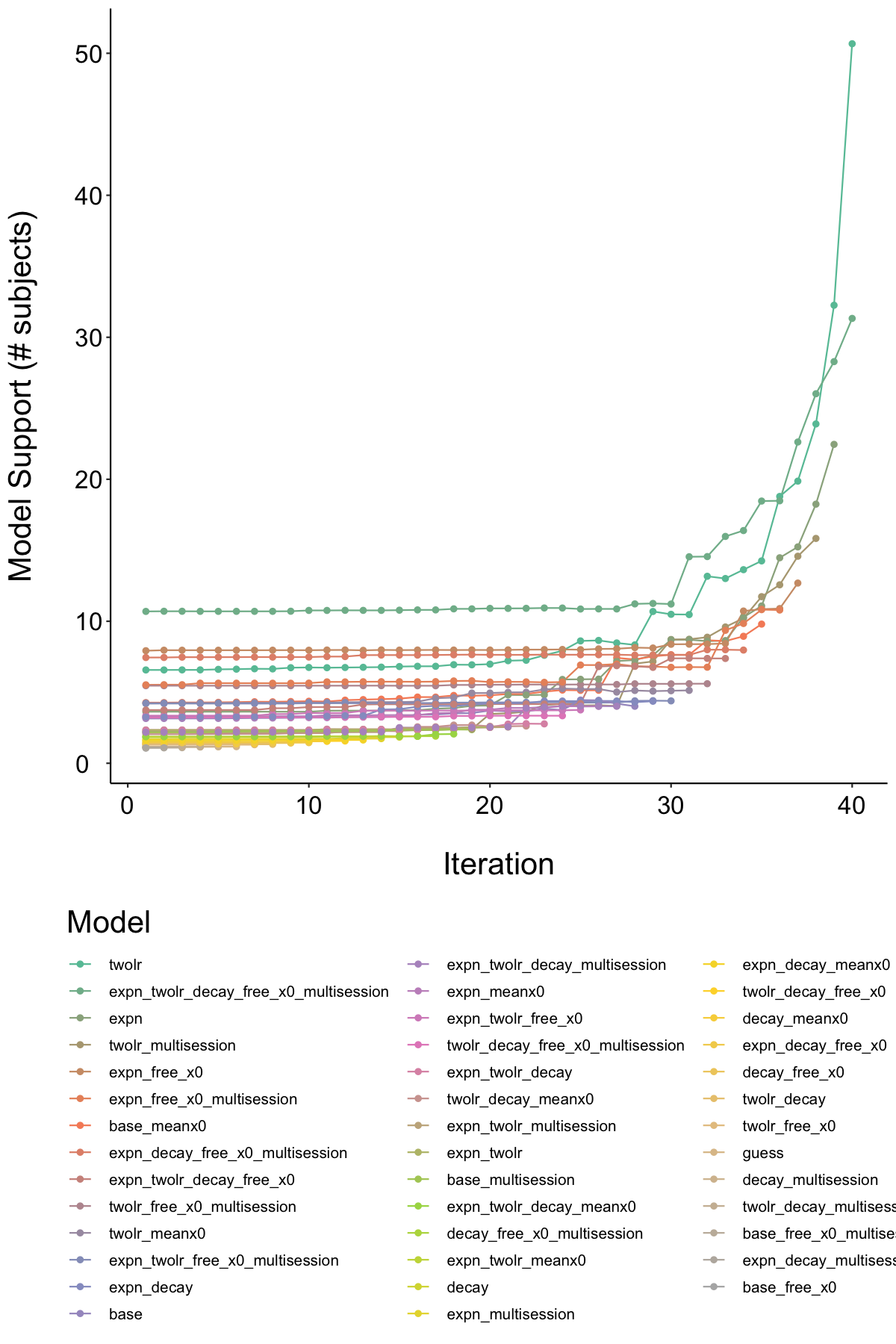

**Supplemental Figure S1.** Iterative model selection based on Variation Bayes Analysis. All models were initially subjected to a Bayesian Model Comparison to estimate total model support across participants. On each iteration, the model with least support was dropped. Legend is sorted by model rank, as determined by iteration of removal.

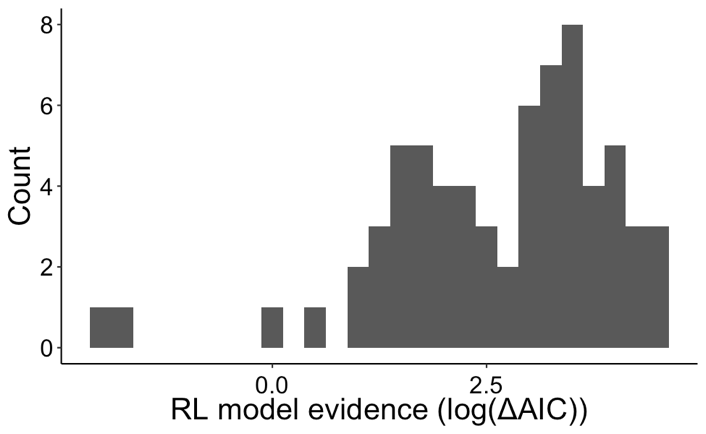

**Supplemental Figure S2.** Distribution of model evidence across subjects, as determined by model AIC relative to a null, “guessing” model in which every choice was accounted for as 50%/50% choice between the two map locations. Greater model support indicates a greater likelihood for the model to be able to predict the actual pattern of choices used by the subject.

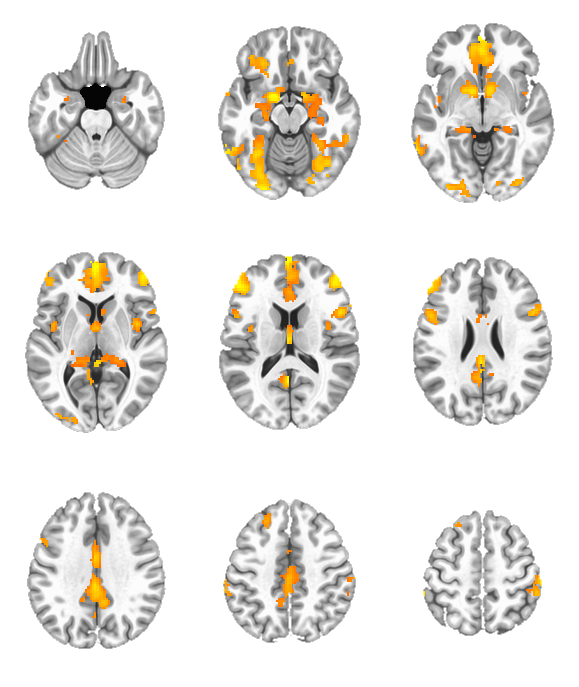

**Supplemental Figure S3.** Activation for the contrast of reward > non-reward trials. Clusters are corrected at 1% FWE (voxel p<0.005, n>96 contiguous voxels), controlling for mean FD at the voxel level. Prominent clusters of activation are seen in the striatum, cingulate cortex, vmPFC, and more (see Supp. Table S1).

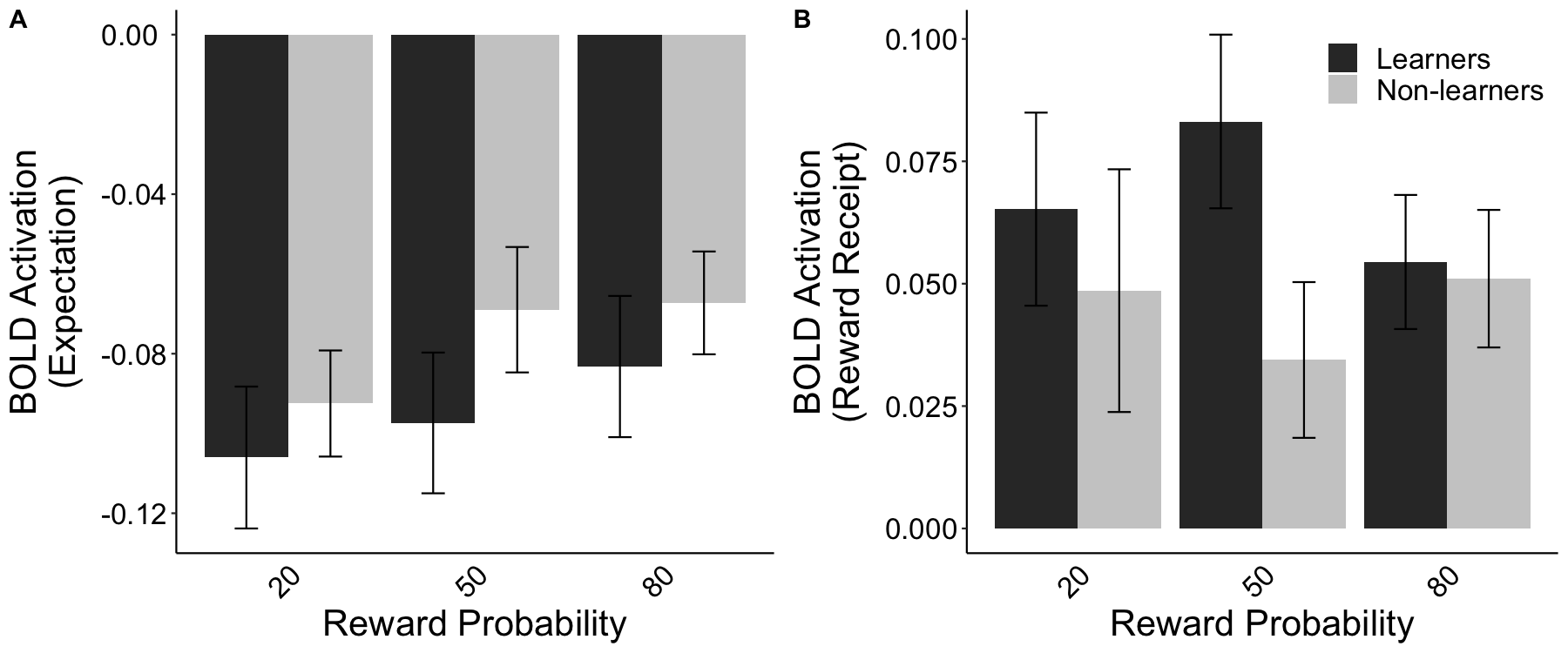

Supplemental Figure S4. BOLD VS activation across task conditions. (A) No group differences in the peak activation in the reward expectation decision-making epoch prior to responding, and (B) Group activation on rewarded trials at the reward receipt epoch. Responses are separated based on the reward probability of the selected map location (20/50/80% probability of a reward).

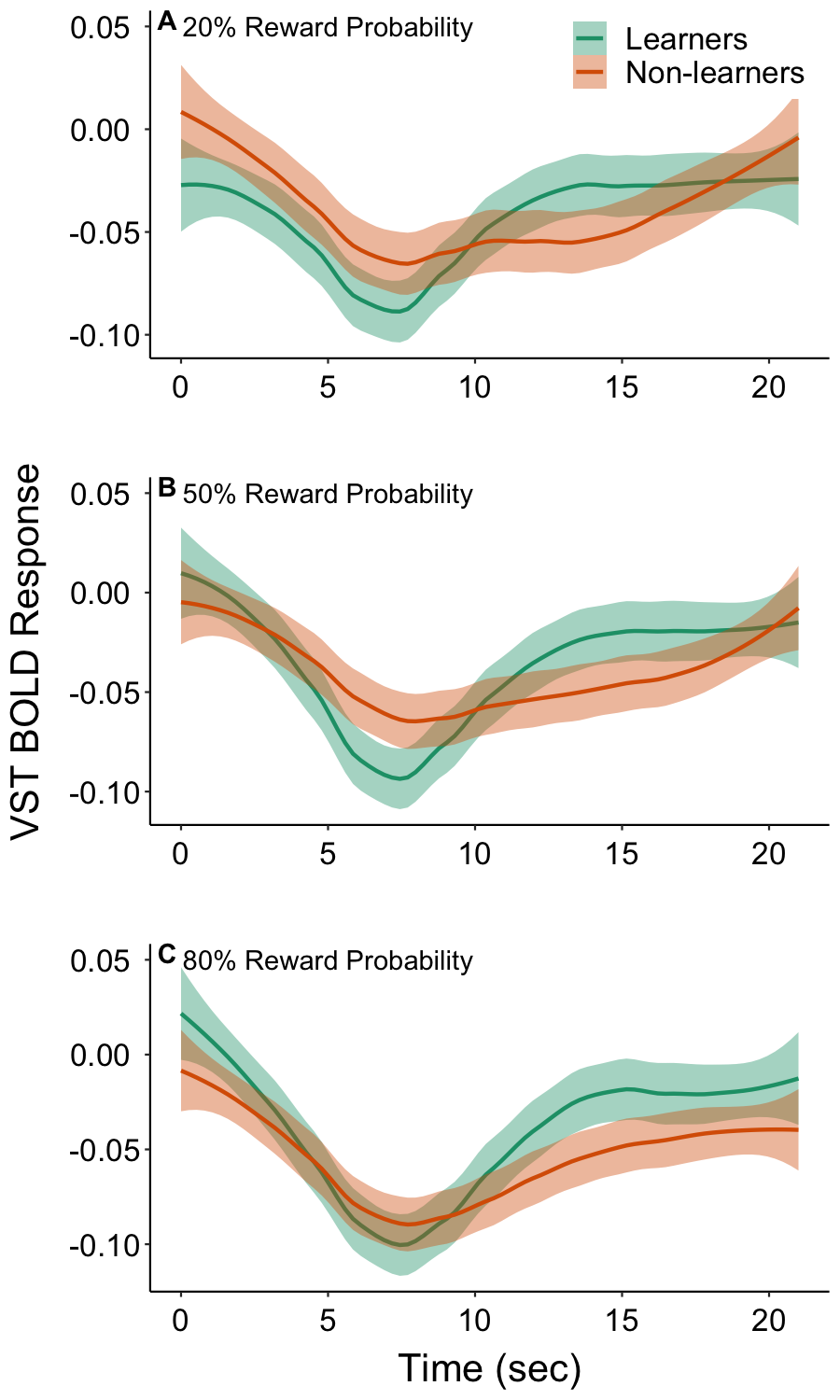

**Supplemental Figure S5.** Timecourses of BOLD response to the expectation epoch of the task. Time is modeled relative to the hash marks appearing indicating the two map locations the subject can choose between on a given trial. Group time courses are shown with LOESS smoothing. Shaded regions indicated +/- 1 SEM.

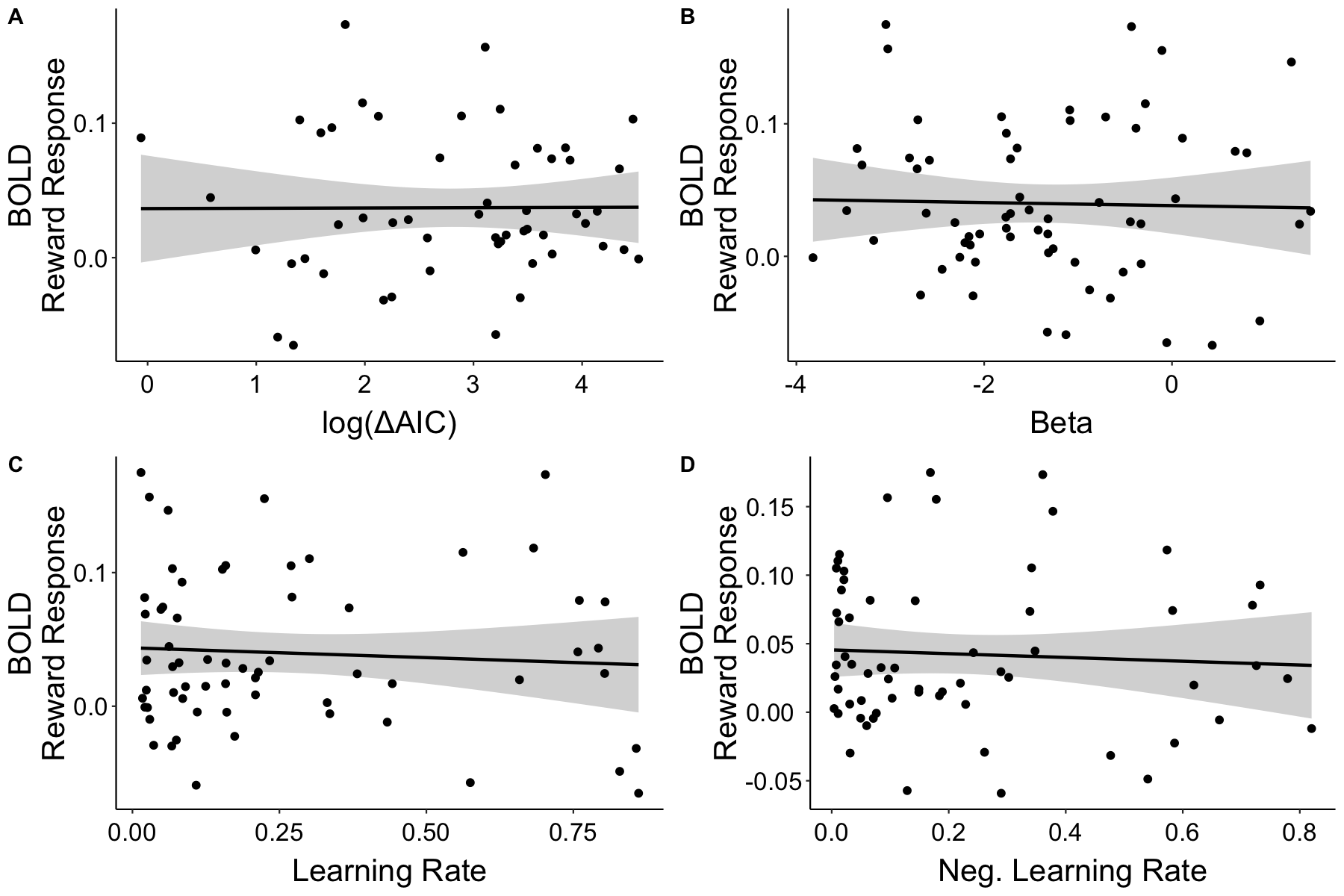

**Supplemental Figure S6.** Correlation between RL model fit parameters and aggregate BOLD reward responses (all rewarded trials vs. all non-rewarded trials), including (A) overall model evidence relative to a guessing model, (B) softmax temperature parameter, (C) learning rate for positive PE trials, and (D) learning rate for negative PE trials.. No significant associations were observed between BOLD reward responses and any of the model parameters.
